# Fishing pressure impacts on anchovy and hake off Peru

**DOI:** 10.64898/2026.09.28.754961

**Authors:** Mariana Hill

## Abstract

The Peruvian anchovy (*Engraulis ringens*) is a small pelagic fish. It is the largest single-species fishery in the world and it is used mainly for the production of fishmeal and fish oil. The Peruvian hake (*Merluccius gayi*) is a predatory demersal fish that is valued for direct human consumption, mainly as frozen products. In this study, I compared the response of both species to certain fishing mortality scenarios using a climatological set-up of the multispecies model OSMOSE for the northern Humboldt Current System. I observed that hake landings benefit from a decreased fishing pressure. In addition, the resilience of the anchovy fishery may increase if a higher threshold for their minimum catch size was implemented. The results of this study provide insights into better management strategies that could be beneficial for the two species based on their life–strategies.

## 1 Introduction

The waters off Peru, in the northern Humboldt Current System (NHCS), are the most productive part of the ocean in terms of fish (Bakun and Weeks, 2008). They host the Peruvian anchovy (*Engraulis ringens*) which is the largest single-species fishery of the planet (Ñiquen Carranza et al., 2000; Chavez et al., 2003). Its production peaked in 1971 at 12.3Mt (Aranda, 2009). It is mainly used to produce fishmeal and fish oil. As the largest producer of these products, Peru generated in average 1.7 and 0.27 Mt of fishmeal and fish oil, respectively, between 2001 and 2006 (Péron, François Mittaine, and Le Gallic, 2010). Fishmeal is used mainly in aquaculture as food and also to feed land stock animals which are eventually consumed by humans (Shepherd and Jackson, 2013). Fish oil is used in aquaculture and for direct human consumption as nutritional supplements (St. John et al., 2016).

Anchovy is a small pelagic fish living in large congregations in the nutrient-rich waters off Peru between 0 and 60 m depth (Ñiquen Carranza et al., 2000). Its main source of food are euphausiids followed by copeopods (Espinoza and Bertrand, 2008). It spawns all year round with the biggest peak between September and November and a smaller peak between February and April. This second peak leads to the greatest recruitment of the year (studies included in Pauly and Tsukayama, 1987). Fishing is prohibited during austral spring and autumn to protect the populations during these spawning peaks. In addition, the minimum catch size of anchovy is 12 cm (Salvatteci and Mendo, 2005; Arias Schreiber, 2012). Anchovy is prone to fluctuations and collapses due to the interannual variability in the oceanographic conditions of the NHCS. It collapsed during the El-Niño events of 1972, 1983 and 1998. It is believed that at least the collapse of 1972 was also influenced by overfishing (Alheit and Niquen, 2004; Arias Schreiber, 2012). Since 1995, the fishery has benefited from regulatory efforts to ensure its sustainability, as well as suitable environmental conditions (Arias Schreiber, 2012).

The Peruvian hake (*Merluccius gayi*) is a large demersal fish that lives in the coastal waters off South America between 1° N and 14° S, extending up to 18° S during El-Niño events (Guevara-Carrasco and Lleonart, 2008). The wider dispersion during this period is thought to have contributed to decreased cannibalism (Guevara-Carrasco and Lleonart, 2008). Hake is valued for human consumption due to its white meat (Del Solar, Sánchez R., and Piazza L., 1965) and it has been industrially exploited since the 1960s to be exported as frozen food (Guevara-Carrasco and Lleonart, 2008). The hake fishery collapsed in 2002 and a moratorium of 20 months was implemented (Guevara-Carrasco and Lleonart, 2008). However, the fishery biomass remained low throughout the 2000s (Guevara-Carrasco and Lleonart, 2008).

Hake and anchovy are two fisheries of the Peruvian upwelling system of high economic importance. They have very different life strategies and both have been subjected to overfishing in the past. In this study, I compared the responses of anchovy and hake to a set if different fishing scenarios. To do so, I employed an end-to-end model of the NHCS ecosystem which simulates nine species of fish and macroinvertebrates, including anchovy and hake. The configuration was calibrated by Hill Cruz et al. (2022) to replicate the average biomasses of anchovy and hake from 2000 to 2008. This study is relevant as a starting point for exploring management strategies that may be successful for each of the two species based on their specific life strategies.

## 2 Methods

For this study, I used the one-way coupled CROCO–BioEBUS–OSMOSE climatological set-up for the NHCS described by Hill Cruz et al. (2022). The physical–biogeochemical component of the system is simulated using the Coastal and Regional Ocean COmmunity model (CROCO Shchepetkin and McWilliams, 2005) coupled to the Biogeochemical model developed for the Eastern Boundary Upwelling Systems (BioEBUS Gutknecht et al., 2013). The higher trophic levels of the system are represented by the Object-oriented Simulator of Marine Ecosystems (OSMOSE Shin and Cury, 2001; Shin and Cury, 2004) which is an individual-based model that simulates the life-cycle of fish. The climatological set-up for the NHCS simulates the mean ecosystem state from years 2000 to 2008. It consists of four plankton groups and nine species of higher trophic levels including the Peruvian anchovy (*Engraulis ringens*) and Peruvian hake (*Merluccius gayi*). Using this configuration as a starting point, I performed 12 simulations changing the fishing rate of anchovy (*F*_*a*_) and hake (*F*_*h*_), as well as the minimum fishing size of anchovy (*A*_*min*_). All experiments were ran for 100 years of spin-up and then 200 years of simulation. When changing the fishing rate of anchovy and hake, I applied two basic scenarios: a) *F*_*i*_*−* means that, during 100 years of spin-up, the fishing rate equals the fishing rate in the control simulation 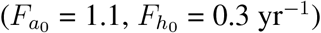, and then the fishing rate is decreased by 5 % of 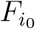 every 10 years until 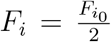 ; for the remaining years 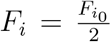. b) *F* + means that the fishing rate of the spin-up also equals 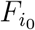 and afterwards 5 % of 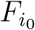 is added to the fishing rate in 10-years intervals. I simulated two sets of experiments. The first set looked at the effects of modifying the minimum fishing size of anchovy, as well as its fishing rate (*A*_*min*_ and *F*_*a*_, respectively). The fishing rate of hake was not modified in these experiments (Table 1). For the second set of experiments, I modified the fishing rates of both hake and anchovy (*F*_*a*_ and *F*_*h*_), without changing the minimum fishing size of anchovy (Table 2). Because OSMOSE is a stochastic model, results vary slightly among replicates of the same set-up. Therefore, I performed each simulation 20 times and averaged the outputs. I reported the running mean of 12 months to filter the seasonal variability.

**Table 1:**
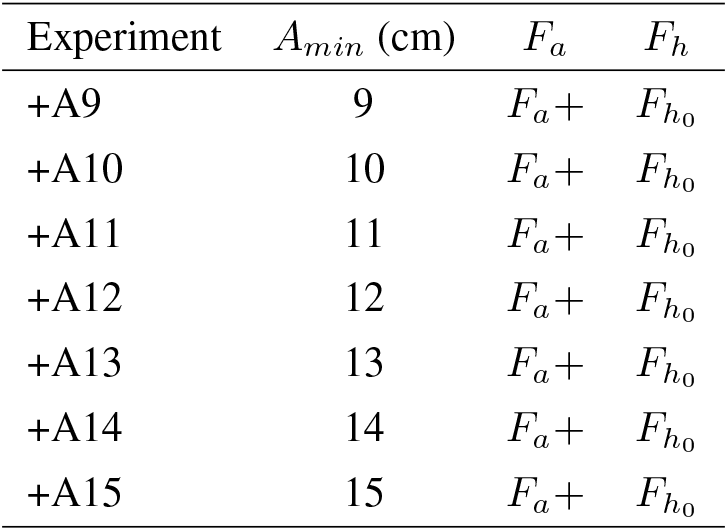
List of experiments changing anchovy minimum fishing size (*A*_*min*_; cm) as well as anchovy fishing rate (*F*_*a*_). The fishing rate of hake is kept at the control value (*F*_*h*_ = 0.3 yr^*−*1^).

| Experiment | $A_{min}$ (cm) | $F_a$ | $F_h$ |
| --- | --- | --- | --- |
| +A9 | 9 | $F_a +$ | $F_{h_0}$ |
| +A10 | 10 | $F_a +$ | $F_{h_0}$ |
| +A11 | 11 | $F_a +$ | $F_{h_0}$ |
| +A12 | 12 | $F_a +$ | $F_{h_0}$ |
| +A13 | 13 | $F_a +$ | $F_{h_0}$ |
| +A14 | 14 | $F_a +$ | $F_{h_0}$ |
| +A15 | 15 | $F_a +$ | $F_{h_0}$ |

**Table 2:** List of experiments changing anchovy and hake fishing rates (*F*_*i*_). The minimum fishing size of anchovy remains constant (*A*_*min*_ = 12 cm).

| Experiment | $A_{min}$ (cm) | $F_a$ | $F_h$ |
| --- | --- | --- | --- |
| control | 12 | $F_{a_0}$ | $F_{h_0}$ |
| +A12 | 12 | $F_a +$ | $F_{h_0}$ |
| -A | 12 | $F_a -$ | $F_{h_0}$ |
| +H | 12 | $F_{a_0}$ | $F_h +$ |
| -H | 12 | $F_{a_0}$ | $F_h -$ |

## 3 Results

First, I examine the effects of increasing and decreasing the minimum catch size of anchovy in addition to increasing its fishing rate. The minimum catch size of 12 cm allows for landings almost as high as 600 Kt per month at the control fishing rate of 1.1 yr^*−*1^ (Figure 1B, black line). There is a tipping point at a fishing rate of about 1.8 yr^*−*1^, around year 125, where the fishing pressure is so high that the landings start to decrease as the fishing rate keeps increasing (Figure 1B, black line). Decreasing the minimum catch size of anchovy switches the tipping point to a fishing pressure almost as low as the 1.1 yr^*−*1^ rate at the beginning of the simulation (Figure 1B, yellow line). Interestingly, decreasing the minimum catch size of anchovy any further than 11 cm does not increase the landings (Figure 1B). Furthermore, it generates a stronger decrease in landings when fishing pressure increases. Increasing the minimum catch size of anchovy results in lower landings at the beginning of the simulation (Figure 1B, blue lines). Nonetheless, it increases the resilience of the species to fishing pressure (Figure 1A, blue lines) enough to allow for a steady increment in catches up to fishing rates as high as 2.2 yr^*−*1^ by year 200 (Figure 1B, blue lines).

**Figure 1:**
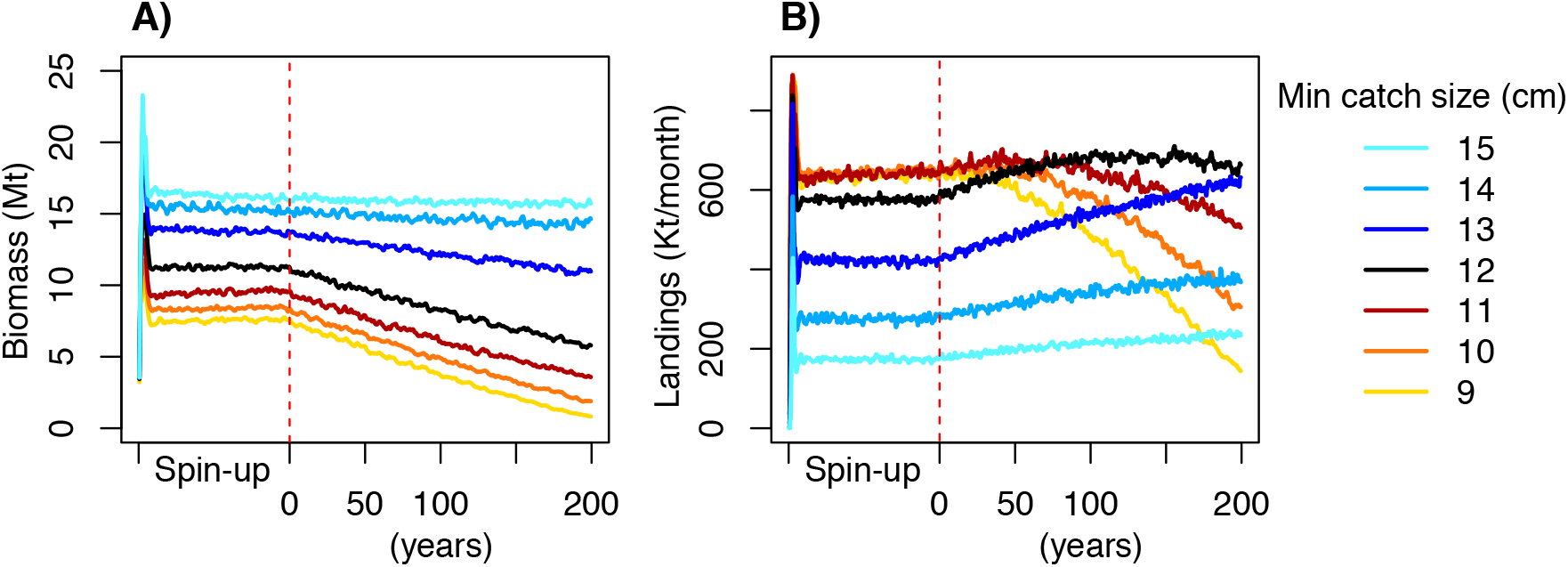
Annual running mean of the anchovy biomass (A) and monthly landings (B) for scenarios +A9 to +A15. Fishing rate is 1.1 yr^*−*1^ during 100 years of spin-up and then it increases by 5 % every ten years for 200 years of simulation. There is a peak close to the beginning of the spin-up because OSMOSE is initialised by releasing eggs during the first 12 years of the simulation (see Hill Cruz et al., 2022).

Hake exhibits a higher sensitivity than anchovy to changes in the fishing rate (Figure 2) and it is driven to a collapse when the fishing rate doubles (Figure 2, +H). Its biomass increases more than twice when the fishing rate is halved (Figure 2, –H). This increase is high enough to generate an increase of about two thirds in the landings, despite the lower fishing rate (Figure 2, –H).

**Figure 2:**
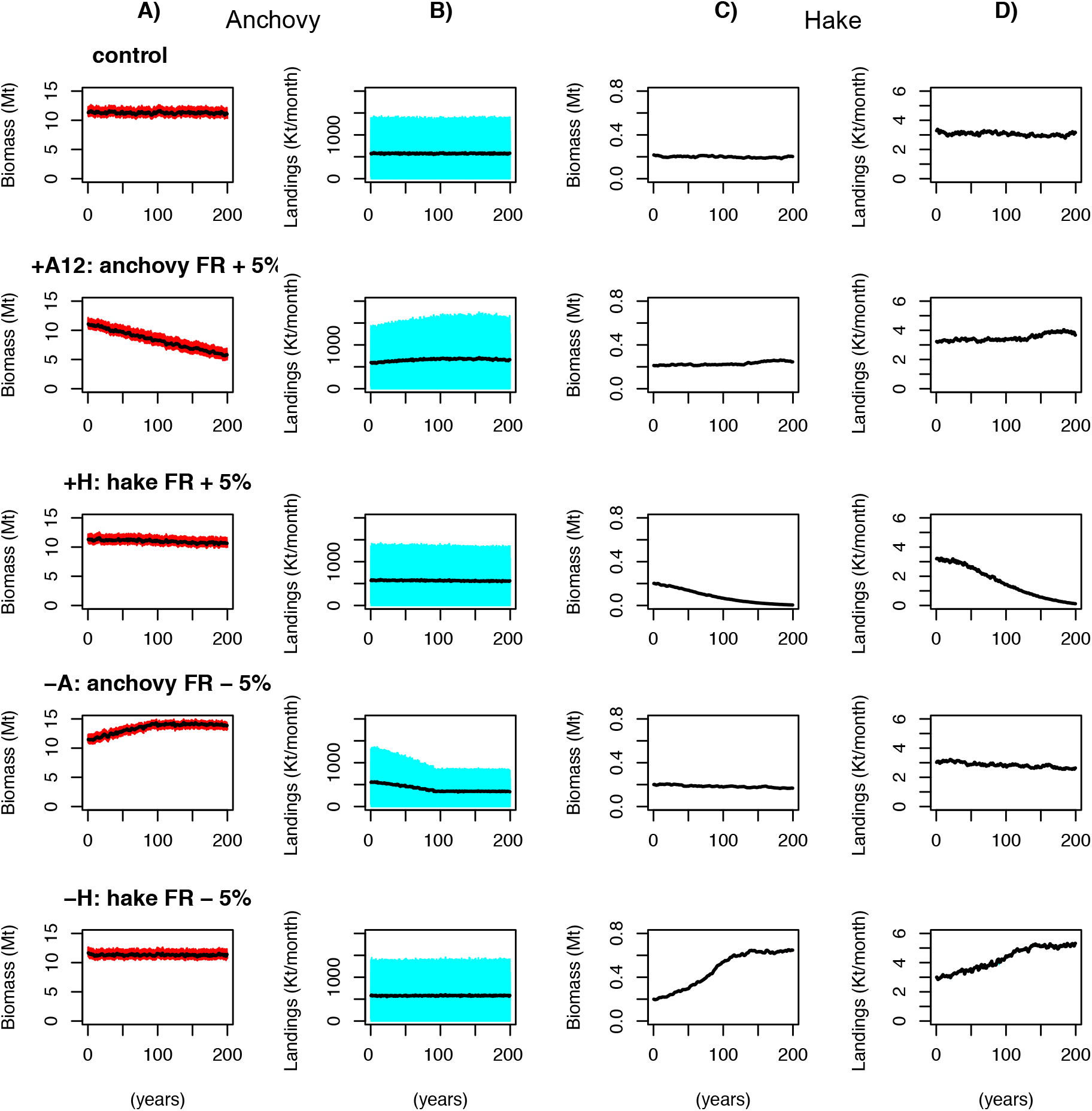
Anchovy (A, B) and hake (C, D) biomass (red) and monthly landings (cyan). Black lines provide the running mean of 12 months. Anchovy fishing is subjected to seasonal fishing closures. Therefore, monthly landings fluctuate between zero and their maximum during the fishing season. Control fishing rates are 1.1 yr^*−*1^ and 0.3 yr^*−*1^ for anchovy and hake, respectively. Fishing rates are constant during spin-up (not shown). After spin-up, fishing rates (FR) are either increased or decreased by 5% every 10 years. See Sect. 2 and Tab. 2 for details on the specific experiments.

In summary, anchovy landings usually increase as fishing rate increase, despite a decrease in its biomass, up to certain threshold. In the case of hake, landings benefit more from a decrease in the fishing rate and are prone to collapse when the fishing rate increases.

## 4 Discussion

This study explored the response of two commercially exploited species, the Peruvian anchovy and the Peruvian hake, in the northern Humboldt Current System (NHCS), to fishing scenarios.

Anchovy is a small pelagic fish that lives up to 4 years (Marzloff et al., 2009) and it is considered to reach maturity at 12 cm (Marzloff et al., 2009). In this study, an increase of fishing pressure in anchovy generates a tipping point at a fishing pressure about two thirds higher than the control. Here I call “tipping point” the point at which any further increase in the fishing pressure does not increase the landings but rather reduces them. Decreasing the minimum catch size does not show a considerable improvement in the landings and makes them rather more vulnerable to a tipping point when the fishing pressure increases. On the other hand, increasing the minimum catch size improves the resilience of the population to a higher fishing pressure. Larger individuals produce more eggs and have longer spawning seasons (Pauly and Soriano, 1987). Hence, this might be an alternative measure to protect the fishery from overfishing, especially after years of recruitment failure. In addition, Salvatteci and Mendo (2005) pointed out that, while the minimum catch size is 12 cm, with a tolerance for individuals smaller than this size of 10 % of the catch, smaller individuals have been harvested. In my study, the minimum catch size was set to 12 cm with no tolerance range and assuming no bycatch. Therefore, the control scenario in this study is rather conservative. Furthermore, Pauly and Soriano (1987) reported a 50 % maturity of individuals at 14 cm. This is further supported by Ñiquen Carranza et al. (1999). Thus, in reality, the 12 cm minimum catch size may be removing a considerable amount of individuals that have not reproduced yet in the real world.

Hake has longer generation times than anchovy and it is less resilient to overfishing. Fernández Ramírez (1987) reported female individuals as old as 9 years off the coast of Peru. In OSMOSE, the maximum, age of 12 years was employed following Marzloff et al. (2009). Before the 1990s, individuals matured at around 2.5 years of age (Guevara-Carrasco and Lleonart, 2008) and 27 to 29 cm in length (Canal Loayza, 1989), with most spawning individuals being more than 3 years old (Guevara-Carrasco and Lleonart, 2008). The age and size at maturity, however, has decreased throughout the years (Guevara-Carrasco and Lleonart, 2008). Large, long–lived species with lower population growth are considered to be less resilient to fishing pressure (Jennings, Greenstreet, and Reynolds, 1999). In this study, I observed that hake is negatively affected by an increased fishing rate to the point of collapsing, (this was not observed for anchovy). On the other hand, decreasing the fishing pressure by half benefits hake by yielding almost 70% higher landings and 200% higher biomass relative to the control means. In contrast, Marzloff et al. (2009) reported an increase in biomass of twice their reference state after a complete removal of the fishing pressure (Marzloff et al., 2009). Both my study and the study by Marzloff et al. (2009) agree that lower fishing pressure would be beneficial for the hake fishery. From this, I conclude, that hake in the model is in an overfished state. The model was calibrated based on the hake state in the 2000s (see Hill Cruz et al., 2022). This period of time corresponds to a known collapse of the hake fishery due to overfishing (Ballón et al., 2008; Guevara-Carrasco and Lleonart, 2008), in agreement with the model. In 2002, a moratorium on fishing hake was implemented for 20 months (Guevara-Carrasco and Lleonart, 2008). This was shorter than the amount of time that takes hakes to reach maturity. Hake population did not show an immediate recovery and hake biomass remained low at least for the following six years (Guevara-Carrasco and Lleonart, 2008).

This study analysed the impacts of increasing fishing pressure on hake and anchovy in an idealised climatological simulation. In reality, the NHCS is affected by interannual variability, specifically El Niño–Southern Oscillation (Barber and Chavez, 1983; Fiedler, 2002; Alheit and Niquen, 2004), as well as regimes of cold and warm water (Chavez et al., 2003; Alheit and Niquen, 2004). During El Niño conditions, anchovies migrate to deeper waters, closer to the coast and further south (Ñiquen Carranza et al., 2000). Major anchovy collapses in the past (e.i., 1972–1973, 1982–1983 as well as the decline in 1997–1998) have been associated with recruitment failure during El Niño events (Boerema and Gulland, 1973; Clark, 1976; Alheit and Niquen, 2004). Therefore, I would expect that an interannual simulation that replicates the environmental effect on anchovy would result in a lower resilience to fishing pressure than the climatological set-up. On the other hand, larger longed–lived fish have been considered to have a higher resilience to environmental variability than smaller fish (MacCall, 2002; Hsieh et al., 2010; Maselko, Andrews, and Hohenlohe, 2020). In the case of hake, it is not yet clear whether the El Niño event has a positive or negative impact. While it stresses the fish due to reduced food availability (Ballón et al., 2008) it also expands its area of occurrence southwards, decreasing cannibalism and catchability (Guevara-Carrasco, Rodríguez, and Rodríguez, 2004). Future studies should consider the impact of fishing pressure in a setting with interannual variability. To do so, it is first necessary to capture the underlying links to the environment that are responsible of the interannual variability in anchovy and hake populations.

In this study, I focused only on anchovy and hake, without looking at the impacts on other species of the ecosystem. However, in recent years, the fisheries management paradigm has been switching from the traditional single–species management to an ecosystem based fisheries management (see Marasco et al., 2007). In this context, the multispecies maximum sustainable yield refers to the maximum yield that can be provided by a system rather than a single fishery (Worm et al., 2009). In the case of the NHCS, the exploitation of small pelagic fish shifted from being monospecific, mainly anchovy, in the 1960s, to multispecific after the anchovy collapse of 1972 and increase in the populations of sardine (*Sardinops sagax*) and jack and chub mackerel (*Trachurus murphyi* and *Scomber japonicus*, respectively) (Ñiquen Carranza et al., 2000). Therefore, it is worth to consider the effect of fishing pressure on these species as well in further studies. Smith et al. (2011) pointed out that reducing the fishing pressure on small fish by half would only decrease the maximum sustainable yield to 80 % while providing large benefits to the surrounding ecosystem. In addition, the potential value, not only yield, of the ecosystem should also be considered (Bieg and McCann, 2020). Only 1 % of the anchovy landings are used for direct human consumption as canned food (Ñiquen Carranza et al., 2000). Hake, on the contrary, is valued for direct human consumption and it is industrially commercialised as frozen food (Guevara-Carrasco and Lleonart, 2008). A management strategy that aims not necessarily to the optimal exploitation of anchovy, but also to rebuilding other species in the region with higher value, might be an alternative to increase the profits while ensuring the health of the ecosystem (see Bieg and McCann, 2020). Finally, rethinking the value of small pelagic fish for direct human consumption might be an alternative to increase the ecosystem profits (Christensen et al., 2014) and also to ensure global food security (Tacon and Metian, 2009).

## 5 Conclusion

In this study, I showed potential outcomes of alternative management scenarios for anchovy and hake. For the simulated years, hake shows to be more sensitive to fishing pressure and a reduced fishing pressure may be beneficial for hake landings. On the other hand, anchovy exhibits a higher resilience to increased fishing rate in this climatological set-up, especially when increasing the minimum catch size. Fishing only adult anchovies is important to ensure the resilience of the population, especially under increased fishing pressure. This study focused merely on the top-down aspect and on two species. Further work should look at the effects of changing fishing management scenarios for all targeted species of the ecosystem in combination to environmental –interannual–, variability.

## Acknowledgements

I would like to thank Miguel Ñiquen and Yunne Shin for inspiring this study. The work presented in this study is part of my doctoral thesis (Hill Cruz, 2022). This work received financial support by the Bundesministerin für Forschung, Technologie und Raumfahrt (BMFTR) through the projects Coastal Upwelling System in a Changing Ocean CUSCO (03F0813 A) and Humboldt-Tipping (01LC1823B). I gratefully acknowledge the computing time granted by the Resource Allocation Board and provided on the supercomputer Lise and Emmy at NHR@ZIB and NHR@Göttingen as part of the NHR infrastructure. Simulations were performed using the computing facilities of NHR.

